# Systematic Chromosome Rearrangement Induced by CRISPR-Cas9 Reshapes the Genome and Transcriptome of Human Cells

**DOI:** 10.1101/2021.06.15.448470

**Authors:** Ying Liu, Guangwei Ma, Zenghong Gao, Jian Li, Jin Wang, Xiangping Zhu, Jiawen Yang, Yiting Zhou, Kaishun Hu, Yin Zhang, Yabin Guo

**Affiliations:** Guangdong Provincial Key Laboratory of Malignant Tumor Epigenetics and Gene Regulation, Medical Research Center, Sun Yat-sen Memorial Hospital, Sun Yat-sen University, Guangzhou, 510120; Ministry of Education Key Laboratory for Ecology of Tropical Islands, Key Laboratory of Tropical Animal and Plant Ecology of Hainan Province, College of Life Sciences, Hainan Normal University, Haikou 571158; State Key Laboratory of Biocontrol, School of Life Sciences, Sun Yat-sen University, Guangzhou, China

**Keywords:** CRISPR-Cas9, chromosome rearrangement, human genome, retrotransposon, LINE-1, Alu

## Abstract

Chromosome rearrangement plays important roles in development, carcinogenesis and evolution. However, its mechanism and subsequent effects are not fully understood. At present, large-scale chromosome rearrangement has been performed in the simple eukaryote, wine yeast, but the relative research in mammalian cells remains at the level of individual chromosome rearrangement due to technical limitations. In this study, we used CRISPR-Cas9 to target the highly repetitive human endogenous retrotransposons, LINE-1 (L1) and Alu, resulting in a large number of DNA double-strand breaks in the chromosomes. While this operation killed the majority of the cells, we eventually obtained live cell groups. Karyotype analysis and genome re-sequencing proved that we have achieved systematic chromosome rearrangement (SCR) in human cells. The copy number variations (CNVs) of the SCR genomes showed typical patterns that observed in tumor genomes. For example, the most frequent deleted region Chr9p21 containing p15 and p16 tumor suppressor, and the amplified region Chr8q24 containing MYC in tumors were all identified in both SCR cells. The ATAC-seq and RNA-seq further revealed that the epigenetic and transcriptomic landscapes were deeply reshaped by the SCR. Gene expressions related to p53 pathway, DNA repair, cell cycle and apoptosis were greatly altered to facilitate the cell survival under the severe stress induced by the large-scale chromosomal breaks. In addition, we found that the cells acquired CRISPR-Cas9 resistance after SCR by interfering with the Cas9 mRNA. Our study provided a new application of CRISPR-Cas9 and a practical approach for SCR in complex mammalian genomes.

**Highlights:** Repetitive retroelements with large copy numbers were targeted using CRISPR-Cas9 in human cells.

Cells survived after their chromosomal DNAs were heavily cleaved and systematic chromosome rearrangement was achieved.

Systematic chromosome rearrangement reshaped the genome and transcriptome of the cells.

## Introduction

Chromosome rearrangements are mutations that cause genomic structural variations, including insertions, deletions, duplications, copy-number variations (CNVs), inversions and translocations. Chromosome rearrangements are usually caused by DNA double strand breaks (DSBs) and rejoins^1^. It is known that 0.5% of neonatal genomes have abnormalities caused by chromosome rearrangements^2^. The famous Robertsonian translocations occur between the human acrocentric chromosomes (chr13, 14, 15, 21, 22 and Y), which may have normal phenotype, but sometime can cause Downs syndrome or other diseases^3^. Chromosome rearrangements also contribute for carcinogenesis^4^. For example, the Philadelphia chromosome is a rearrangement between chromosome 9 and 22, which makes ABL1 and the strong promoter of BCR (break point cluster region) fuse to form BCR-ABL chimeric gene, resulting in continuous high expression of ABL1 kinase and cell transformation^5^. Moreover, Chromosome rearrangement is a major motivation for evolution^6^. Chromosome rearrangements not only change the primary structure of DNA, but also change the three-dimensional (3D) conformation of chromatins. One of the most extreme examples so far is that Shao et al. Combined 16 chromosomes of *Saccharomyces cerevisiae* into one, creating a yeast strain with only one chromosome in 2018, and later they further cyclized this large chromosome into a circular chromosome, like a typical prokaryotic chromosome^9^. This process greatly changed the 3D structure of the yeast genome. But surprisingly, the growth of this strain (with singular linear chromosome) is similar to that of the wild type^8^. Of course, its gene expression profile has changed significantly. Also in yeast, Jef Boeke team developed a method called SCRaMbLE, in which loxP sites are inserted into synthetic chromosomes and Cre recombinase is used to trigger chromosome rearrangements^10–12^. Many interesting results have been obtained using this method, which provides a great facility for the study of chromosome structure (including 3D structure) and function^13^. As multicellular organisms face many more challenges, such as cell differentiation, development and homeostasis, the mammalian genomes are more sophisticatedly regulated at both 3D structure and epigenetics level. Obviously, if similar studies can be performed in the mammalian cells, it would be more helpful for us to understand the roles of chromosome structure in the development and diseases. However, mammalian genomes are far larger and more complex than the yeast genome and a systematic method to induce chromosome rearrangements is yet absent, which limits the research in this field. In recent years, CRISPR-Cas9 genome editing technology has made great progress. Under the guidance of sgRNA, Cas9 endonuclease specifically cleaves DNA sequences, and the cleaved DNA strands were subsequently rejoined via non-homologous end joining (NHEJ) pathway^14,15^. There are large numbers of repetitive sequences in the mammalian genomes^16^. If sgRNAs are designed according these sequences, a large number of chromosome breaks can be generated, which will lead to systematic chromosome rearrangements (SCR) (Fig. 1A). Here, we developed a method called <u>C</u>hromosome <u>Rea</u>rrangement by <u>C</u>RISPR-Cas9 (CReaC). We used sgRNAs to target endogenous retrotransposons, LINE-1 (L1) or Alu in HEK293T cells and obtained cells with significant different karyotypes from the control cells. Whole genome sequencing (WGS) showed that large numbers of inversions, translocations and CNVs have occurred. Then we further performed transcriptomic and epigenetic studies to evaluate the effect of the SCR on the cell physiological status.

**Figure 1.**
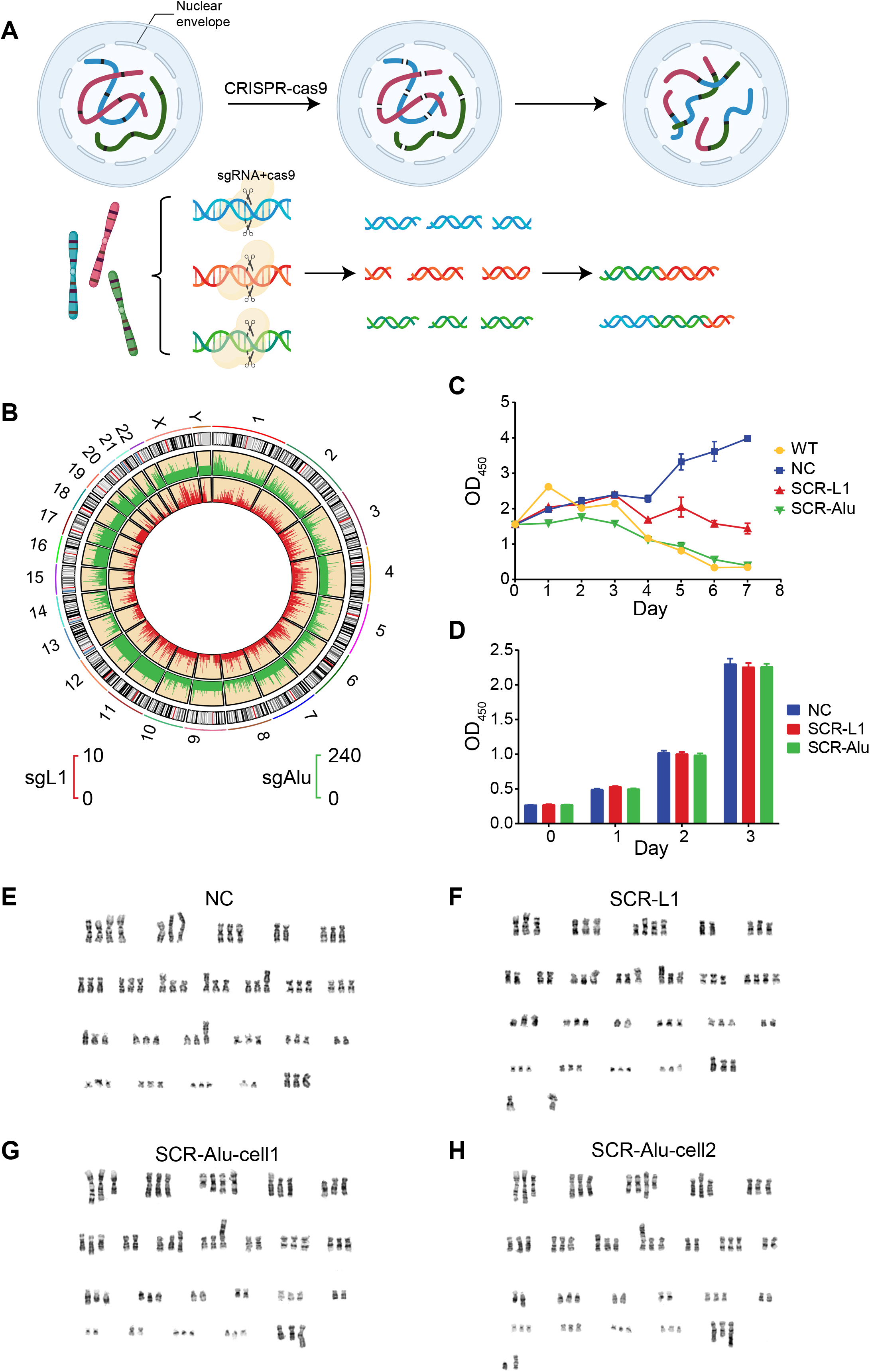
The design of CReaC, cell selections and cytogenetic assay. A, the mechanism that CReaC works. B, the distributions of sgL1 and sgAlu on the human chromosomes, the red bars of the inner track is for sgL1 and green bars of the outer track is for sgAlu. C, the selection if the transfected cells in the presence of puromycin. D, The growth rate of the cells after selection. E-H, karyotype of NC (E), SCR-L1 (F) and SCR-Alu (G & H).

## Results

### Experiment design and cell selection

There are >500000 LINE-1 (intact elements and fragments) and >1000000 Alu copies in the human genome^16^, which provides an ideal set of targets for making multiple DNA DSBs using CRISPR-Cas9. We designed a sgRNA according to the ORF2 region of L1 and a sgRNA according to the most conservative region of Alu, which have 7398 and 317924 matching sites respectively in the current hg38 human genome haploid version (Fig. 1B). These sgRNAs were cloned into the pSB-CRISPR plasmid as previously described and a 20 nt sequences with no match in the human genome were used as negative control sgRNA. The plasmids (pSB-CRISPR-sgNC, pSB-CRISPR-sgL1 and pSB-CRISPR-sgAlu) were transfected together with the SB100X plasmids^18^ into HEK293T cells respectively. The cells were then kept in the presence of puromycin, forcing the Cas9 endonuclease and sgRNAs to express constantly and keep cleaving the chromosomes. The cells that survived after puromycin selection were designated as SCR-L1, SCR-Alu and NC respectively.

As expected, the survival rates of SCR-L1 and SCR-Alu are lower than that of NC, due to the strong stress caused by multiple DSBs (Fig. 1C). The SCR-L1 and SCR-Alu cells recovered, and eventually grew up after roughly three weeks. Then, they started to proliferate stably, and there was no difference between the growth rates of the SCR and NC cells. Thus, new immortalized strains were created by targeting repetitive retroelements with CRISPR-Cas9 (Fig. 1D).

To see if chromosome rearrangements really happened, cytogenetics assay was performed. SCR and NC cells were treated with the classical Giemsa Staining (Fig. 1E-H). It is known that HEK293 cells are female human cells with a karyotype near triploid^19^. Typical NC cells have 67 chromosomes as expected, while SCR cells showed apparent chromosome aberrations, and unrecognizable chromosomes were observed in both of them (Fig. 1F and H). Moreover, one of the SCR-Alu cells contains only 63 chromosomes (Fig. 1G), indicating that CReaC might be used as a tool for genome minimization.

### SCR-L1 and SCR-Alu cells were evaluated using multi-omics approaches

To evaluate the new selected strains comprehensively, we performed WGS, RNA-seq and ATAC-seq (Assay for Transposase-Accessible Chromatin with high throughput sequencing) for the SCR and NC cells (Fig. 2A and B).

**Figure 2.**
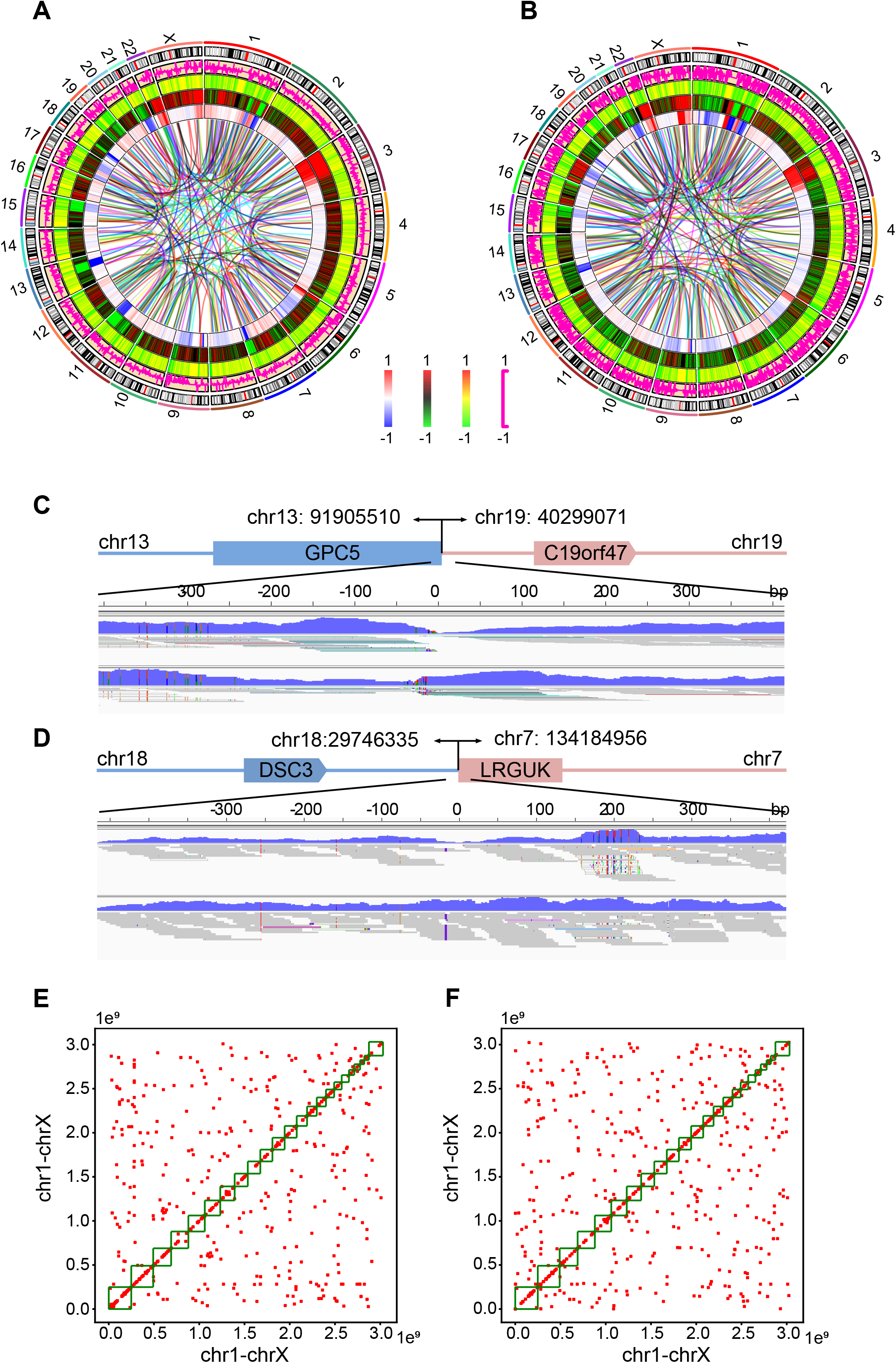
Multi-omics analyses were performed to the SCR and NC cells. A & B, the chromosomal structure, chromatin accessibility and gene expression changes of SCR-L1 vs NC (A) and SCR-Alu vs NC (B). The tracks of the Circos charts from the center to the outer are translocation/inversion, difference of CNV, difference of raw sequence match of ATAC-seq, difference of ATAC-seq peaks and the difference of gene expression. The interval for the heatmaps and curves is 500 kb-interval. C & D, Examples of IGV images for the translocations of SCR-L1 (C) and SCR-Alu (D), the junctions were not covered by sequence reads of WGS in NC, but were covered in SCR cells. E & F, 2D plots show the intra- and interchromosome translocations in SCR-L1 (E) and SCR-Alu (F).

The WGS shows that there were 406 chromosome translocations and inversions took place in SCR-L1 cells, and 495 translocaitons/inversions in SCR-Alu cells (Fig. 2A and B). Two translocation examples of SCR-L1 and SCR-Alu were shown respectively in Fig. 2C and D. This result confirmed that large scale chromosome rearrangements were successfully induced by cleaving DNA double strands at multiple positions with CRISPR-Cas9. Interestingly, although the sites of sgAlu are ~40 times as those of sgL1, the translocation/inversion number of SCR-Alu is very similar to that of SCR-L1. Perhaps, the same selection condition led to the convergence of the two groups of cells.

To see if there are differences between the frequencies of interchromosome translocations and intrachromosome translocations (including inversions), we plotted all the rearrangements events on 2-dimension charts (Fig. 2E and F). Lots of points clustered around the diagonals, indicating that the rearrangements between neighboring retroelements are more common than between retroelements that are far from each other or in different chromosomes. Except for the neighboring rearrangements, there are no difference between the interchromosome rearrangements and the intrachromosome rearrangements. The Hi-C studies showed that there are significantly more intrachromosome interactions than interchromosome interactions^20^. Therefore, the chromosome rearrangements may have little association with the interactions between different chromosomal regions.

RNA-seq showed that the gene expression profiles of both the SCR cells were significantly different from that of NC. And apparently, the change of the expression profile of SCR-Alu is far more drastic than that of SCR-L1 (Fig. 2A and B, pink curves).

The ATAC-seq shows that although the raw sequence reads that matched to the chromosomes were similar between the three groups (Fig. 2A and B, green-black-red heatmap), the peak densities of both SCR strains were dramatically decreased compared to that of NC (green-yellow-red heatmap).

An overview of multi-omics showed that the SCR cells differ from the NC cells in multiple dimensions, including the genomic primary structure, the epigenetic modification and the gene expression profile. Detailed analyses were carried out in the following sections.

### The genomic structure variations of SCR cells

In the Circos chart of SCR-Alu (Fig. 2B), it seems that regions that contain more translocation/inversion events also contain more CNVs. To test this assumption, we compared the numbers of translocation/inversion events in the regions containing high-level CNVs to those in the randomized genomic regions (the shuffled regions of same length as those for evaluating CNVs, see Methods). As shown in Fig. 3A and B, for SCR-Alu, the mean translocation/inversion events in randomized genomic regions is 81, while there are significantly more translocation/inversion events in the regions with high-level CNVs (126 events, P=0.00042). However, similar difference was not found between the regions with high-level CNVs and the random regions in SCR-L1 (P=0.1138), which might be because L1 is much longer than Alu and the subsequent recombination process are more complicated.

**Figure 3.**
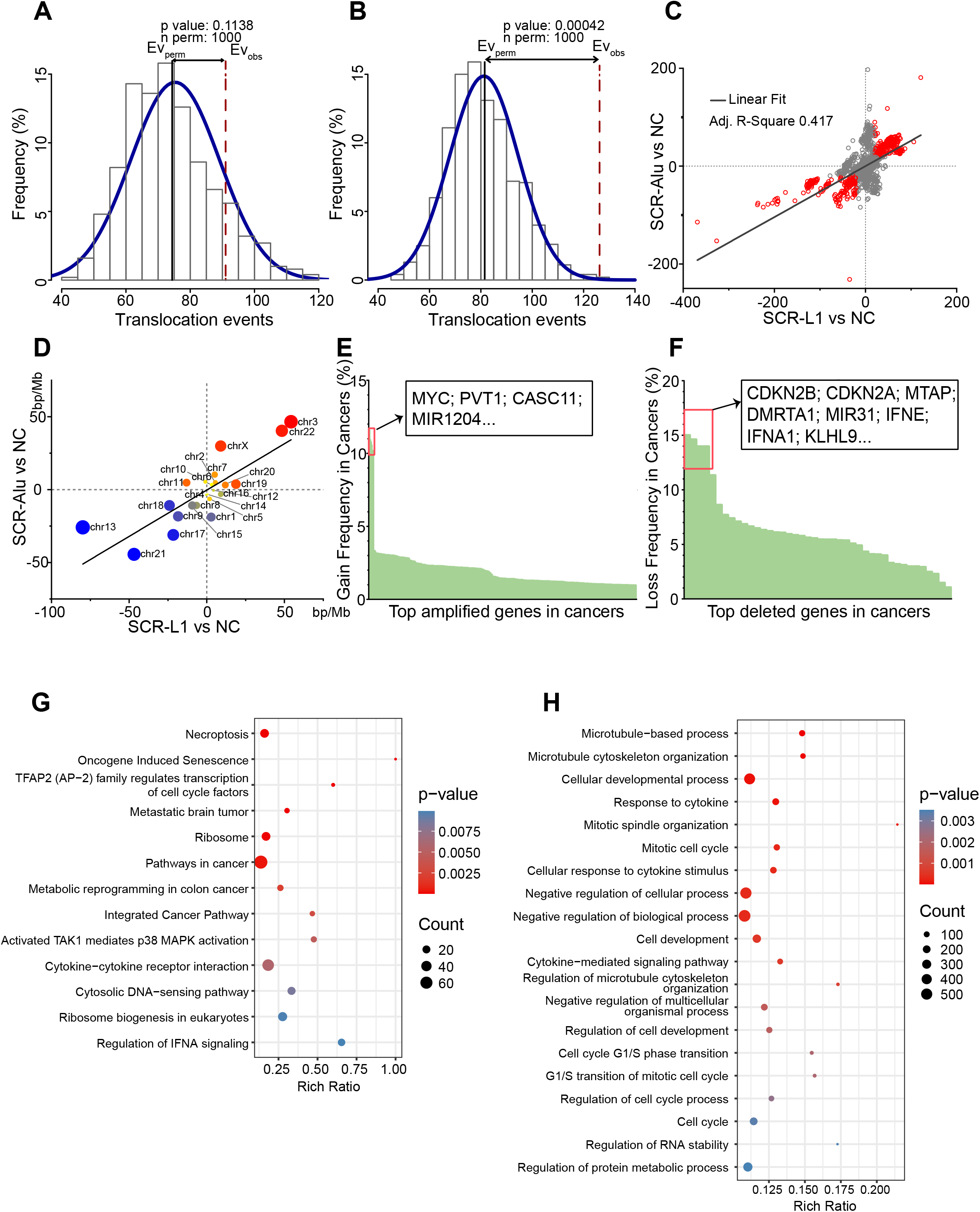
The copy number variations of the SCR cells. A & B, the translocation events in the high CNV regions of SCR-L1 vs NC (A) and SCR-Alu vs NC (B), the genomic regions of 500 kb (same size for calculating CNV) were permutated 1000 times to assess the distribution of translation events of random region. Normal distribution was used to calculate the P value of observed translations in the high CNV regions. C, the correlation the CNVs/500 kb-interval between SCR-L1 vs NC and SCR-Alu vs NC, the red points indicate sequence read matches significantly changed with same trend in the two SCR cells. D, The correlation of genomic gain and loss at the chromosome level between SCR-L1 vs NC and SCR-Alu vs NC. E & F, the gain (E) and loss (F) frequency in clinical tumor samples of the genes at the high CNV regions shared by SCR-Alu vs NC and SCR-L1 vs NC, which was calculated using TCGA clinical samples data obtained from Cosmic. G & H, the high CNVs shared by SCR-L1 vs NC and SCR-Alu vs NC were analyzed using WikiPathway, Reactome, KEGG (G) and GO (H) enrichments.

It seems that the CNV distributions are quite similar between SCR-L1 and SCR-Alu from the Circos chart (Fig. 2A and B), so we tested the correlation between these two distributions based on the same 500 kb intervals as the Circos chart. Fig. 3C showed that the R^2^ between the two distributions was 0.417, indicating a significant correlation. Moreover, if viewed from the chromosome level, the correlation was even stronger (R^2^=0.683, Fig. 2D), which indicated that the CNVs played important roles for the survival of the cells under strong stress. The slopes of the two trend lines also showed that the CNVs in SCR-L1 are more intense than those in SCR-Alu (Fig 3C and D).

Notably, the common CNVs shared by SCR-Alu and SCR-L1 resembled many features of copy number changes that observed in clinical tumor samples, including the most frequent and important regions of gain and loss (Fig. 3E and F). For example, the chr9p21 is the most frequent deleted region in cancer genomes, which contains the important tumor suppressors such as CDKN2A (p16), CDKN2B (p15). p15 and p16 are cyclin-dependent kinase (CDK) inhibitors, that maintain the active state of Rb family members, and promote their binding to E2F1, leading to G1 cell cycle arrest^22^. The loss of p15 and p16 helps the tumor cells bypassing the G1 cell cycle arrest. Moreover, previous study revealed that p16 is required for the reduction in CDK4- and CDK6-mediated Rb kinase activity upon DNA damage^23^. Since the NHEJ repair mainly happens during G1 phase^24^, the simultaneous deletion of Chr9p21 in both SCR cell groups suggests that the loss of p16 and p15 may be a key step for the cells to survive the G1 arrest induced by the multiple NHEJ repairs. Besides, we found the most common amplification of Chr8q24 in tumors also occurred in SCR-L1 and SCR-Alu genomes. This region contains the most famous oncogene, MYC, as well as the onco-lncRNA, PVT1. It is reported that deregulated c-Myc disables the p53-mediated DNA damage response and helps cells with damaged genomes to bypass cell arrest and enter the cell cycle^25^.

The genes in the high-level CNV regions were analyzed using KEGG, Reactome, WikiPathway and Gene Ontology (GO) enrichments. Fig. 3G and H showed that typical pathways related to cell survival such as the “cell cycle”, “G1/S transition” and “pathways in cancer” were enriched, which also indicated the contributions of the CNVs to the survival of the SCR cells.

### Large-scale chromosome rearrangements reshaped the landscape of gene expression

The gene expressions of the SCR cells were compared with the NC cells (Fig. 4A-F; Supplementary Fig. 1C-F). Similar to the sequencing read coverage showed in Fig. 2A and B, both of the gene expressions of SCR-L1 and SCR-Alu were changed greatly. There were 913 mRNAs upregulated and 1396 downregulated in SCR-L1, and 2663 mRNAs upregulated and 1959 downregulated in SCR-Alu (Fig. 4C and D). Similarly, considerable numbers of lncRNAs changed as well (Fig. 4E and F). Obviously, the change of gene expression is much greater in SCR-Alu than that in SCR-L1, which is also consistent to the RNA-seq read coverage (Fig. 2A and B; Supplementary Fig. 1A). Notably, the important onco-lncRNA, MALAT1, was upregulated in SCR-Alu (Fig. 4F; Supplementary Fig. 1F), indicating common cancerogenesis pathways were activated in the SCR cells.

**Figure 4.**
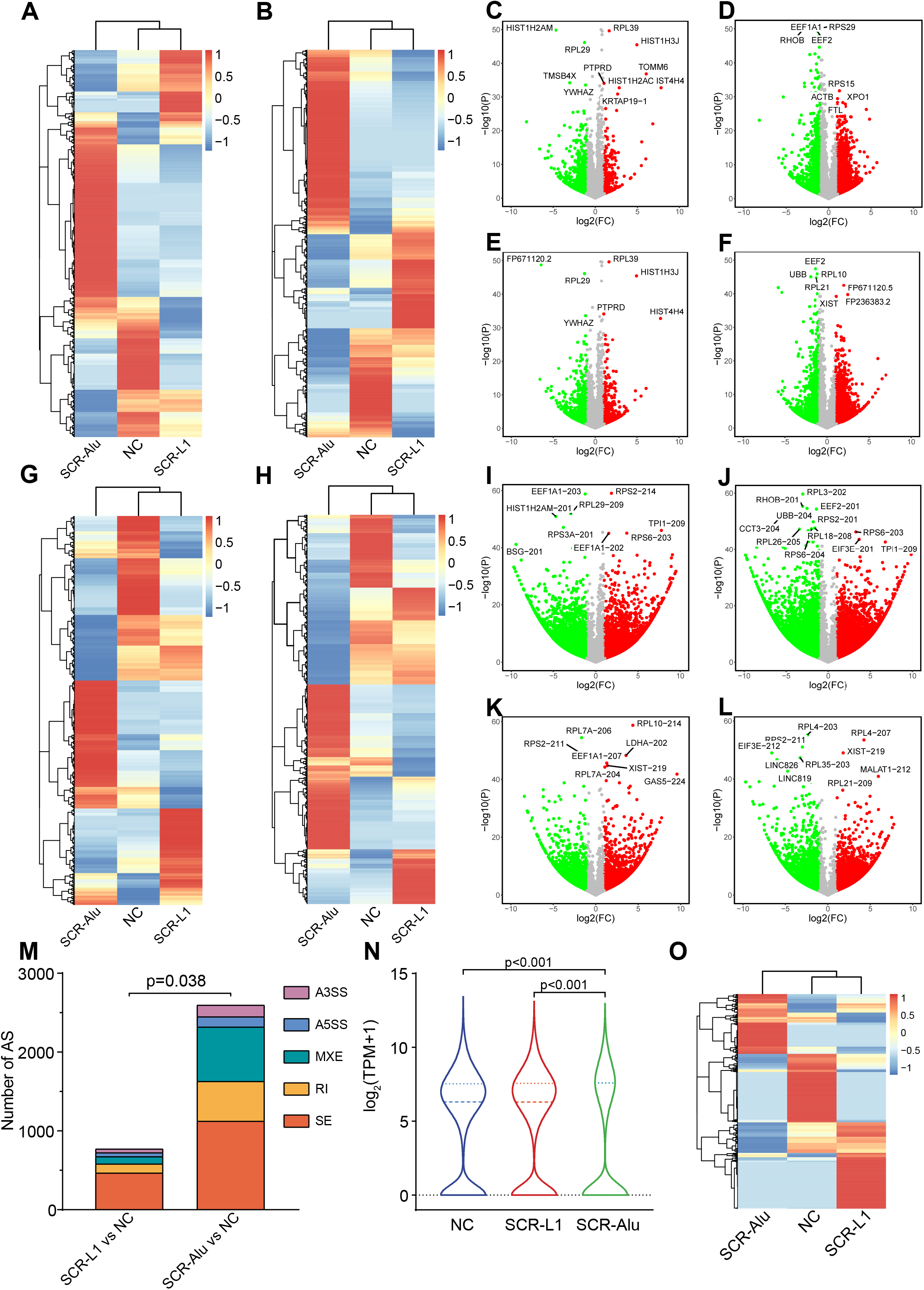
The differential expressed genes (DEGs) between SCR and NC cells. A & B, H-cluster of expression at gene level of mRNAs (A) and lncRNAs (B). C-F, volcano plots show the differential expressions of mRNA (C & D) and lncRNA (E & F) at gene level in SCR-L1 vs NC (C & E) and SCR-Alu (D & F). G & H, the H-cluster of expression at transcript level of mRNAs (G) and lncRNAs (H). I-L, volcano plots show the differential expressions of mRNA (I & J) and lncRNA (K & L) at transcript level in SCR-L1 vs NC (I & K) and SCR-Alu (J & L). M, the different types of alternative splicing of SCR-L1 and SCR-Alu relative to NC. N, violin plot shows the expression of distribution of NC and SCR cells. O, H-cluster of circRNA expressions in NC and SCR cells.

Compared to the expression changes in gene level, the expression changes in transcript level are even more drastic (Fig. 4G-L; Supplementary Fig. 1G-J). The different types of alternative splicing (AS) were shown in Fig. 4M. The AS number of SCR-Alu vs NC is significantly greater than that of ACR-L1 vs NC. It is reported that both L1 and Alu contribute for AS and it seems that the function of Alu on AS is even stronger that that of L1^26–29^. AS of RNA plays important roles in the gene expression regulation of eukaryotes^30^ and aberrant AS is an important contribution for carcinogenesis^31^. Therefore, AS may be a major contribution for the SCR cells to survive the severe stress.

Cluster analysis showed that the gene expression profile of SCR-L1 is relatively close to that of NC, while the gene expression profile of SCR-Alu is far more different (Fig. 4A and B, G and H). The Pearson correlation of the gene expressions also showed the similar result (Supplementary Fig. 1 A and B), i.e., targeting Alu caused substantially larger change than targeting L1 with CRISPR-Cas9.

Besides the expressions of mRNAs and lncRNAs, we also examined the expression of circular RNAs (circRNAs). The most significant change of circRNAs in SCR cells was the decrease of the abundance. Especially in the SCR-Alu cells, the abundance of circRNA is only ~2/3 of that in the NC cells (Fig. 4N). It is known that Alu elements are important for the formation of circRNAs. The homologous sequences of Alu at the flanking introns help the RNA molecules cyclize into circRNAs^32,33^.

The circRNA expressions between cell groups vary greatly. The H-cluster analysis showed even more remarkable contrast than those of the mRNA or lncRNA expressions (Fig. 4O). The differential expressions of circRNAs were shown by volcano plots (Supplementary Fig. 1K and L). Unlike the volcano plots of the mRNAs or lncRNAs, the volcano plots of circRNAs were not symmetric by the line of x=0, rather, the plots shifted leftward, which were also due to the circRNA abundances in SCR-L1 and SCR-Alu were lower than that of the NC. The changes in SCR-L1 and SCR-Alu showed a fairly large overlap (Supplementary Fig. 2A). The KEGG enrichment of the host gene of differentially expressed circRNAs in the two groups also shared multiple pathways. Protein processing in endoplasmic reticulum, ubiquitin mediated proteolysis, RNA transport and cell cycle are the most significant enriched pathways, which also points to the degradation of the abnormal proteins and the response to stress (Supplementary Fig. 2B and C).

### Multiple pathways associated to cell survival were altered in SCR cells

Although SCR-Alu showed very different gene expression profile from SCR-L1, the two SCR strains still shared a lot of common changes, which might be account for the cell survival through severe stress. Fig. 5A showed the log_2_FC of genes from SCR-L1 and SCR-Alu in a scatter plot. The points in the first and third quadrants are the genes that changed with same trend in the two groups of cells. Then the genes with a change fold >2 (log_2_FC > 1 or log_2_FC < −1, the red points in Fig. 5A) and FPKM > 1 were extracted, resulting in a set of 192 genes (Fig. 5B, venn diagram).

**Figure 5.**
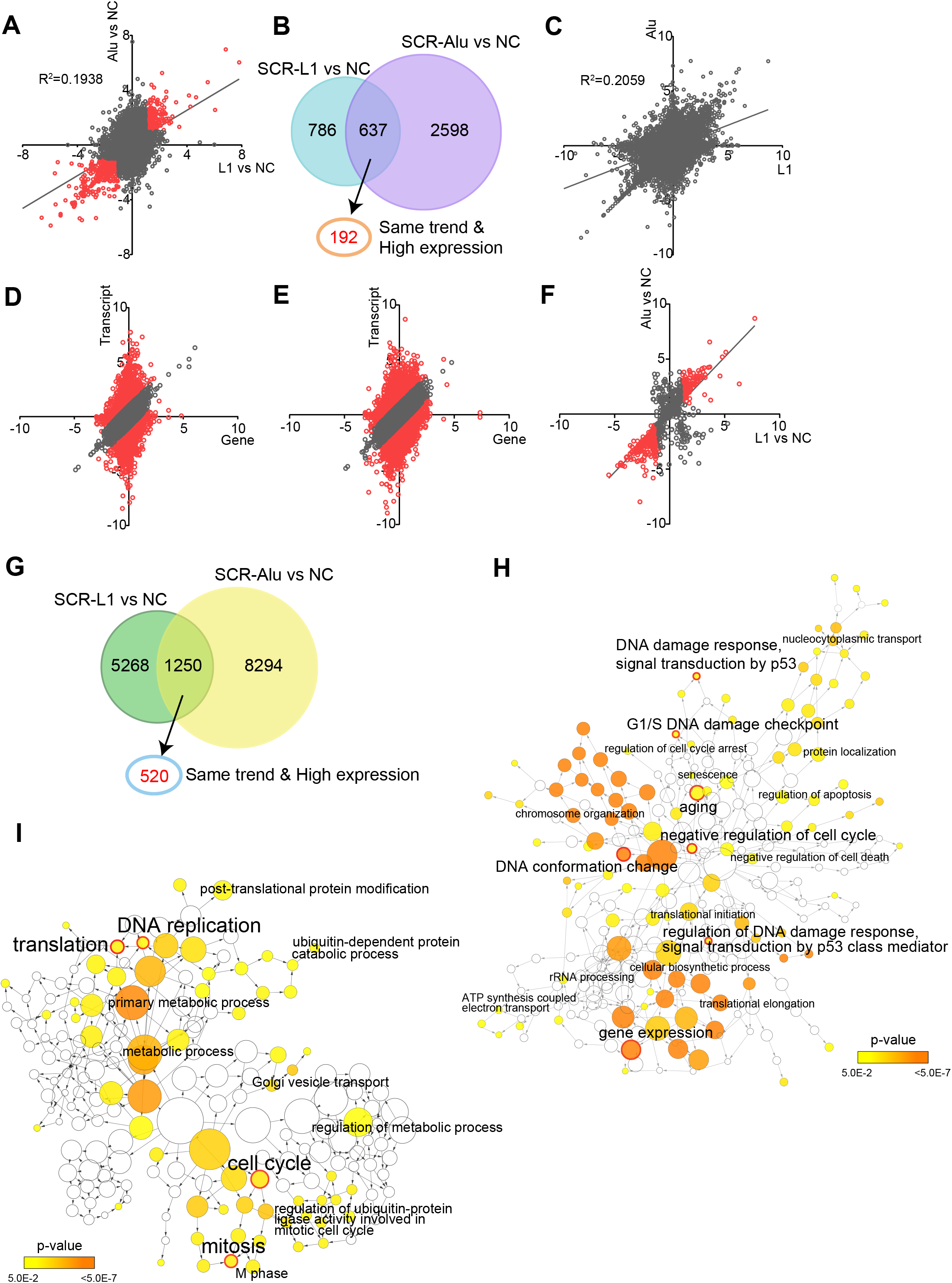
The common expression changes between SCR-L1 and SCR-Alu cells. A, the correlation of expression fold changes at gene level between SCR-L1 vs NC and SCR-Alu vs NC (R2=0.1938), the red points indicate the genes whose expression changed with same trend in the two SCR cells and with fold change ≥ 2. B, the numbers of the genes that marked red in panel A. C, the correlation of expression fold changes at transcript level between SCR-L1 vs NC and SCR-Alu vs NC (R^2^=0.2059). D & E, the expression fold changes at transcript level and gene level in SCR-L1 vs NC (D) and SCR-Alu vs NC (E), red points indicate the genes with fold change at transcript level at least as two folds of the fold change at gene level. F, the correlation between the genes marked red in panel D and panel E, the red points indicate the genes whose expression changed with same trend in the two SCR cells and with fold change ≥ 2. G, the numbers of the genes that marked in panel F. H & I, GO enrichment analysis at gene level (H) and transcript level (I) shown as an interaction network using Cytoscape plug-in, BinGO.

Since the expression changes at transcript level are more significant than those at gene level in the SCR cells, we also examined the common transcript changes in the SCR-L1 and SCR-Alu cells. The correlation of the transcript expressions between the two SCR cells is slightly stronger than that of the gene expressions (Fig. 5C). Next we focus on those genes whose expression changes at transcript level are ≥ 2 folds of the expression changes at gene level (red points in Fig. 5D and E). The transcript changes of these genes showed a fairly strong correlation with an R^2^ of 0.5779 between the two SCR cell groups (Fig. 5F), indicating that the changes of AS were more consistent than those of the overall gene expressions. Therefore, the changes in AS might contribute more than the overall changes of gene expression to the cell survival under stress. Similar to the analysis in Fig. 5B, we still choose the genes with Log_2_FC > 1 at transcript level and with FPKM > 1, and a set of 520 genes (Fig. 5G) were obtained.

We then performed the functional enrichment analysis using GO (Fig. 5H and I), as well as Reactome and WikiPathway (Supplementary Fig. 3) for the genes selected in Fig 5B and G. Notably, almost all the fundamental cellular processes, the DNA synthesis (replication), the RNA synthesis (transcription) and the protein synthesis (translation initiation and extension), were altered in the SCR cells. The most significant enrichment pathways were p53 and DNA damage repair, cell cycle and mitotic check points, apoptosis and some cancer related pathways, which are closely related to the cell survival under stress. The ubiquitination pathway is the major protein post translation modification, which should be due to the large quantity of defect proteins brought by the chromosome rearrangements. This result indicates that SCR has a great physiological impact on cells, so that the cells have to modify almost all the fundamental pathways to survive the crisis.

The p53 pathway is particularly important for the cells upon DNA damage, and it is reported that cells with TP53 mutations are more likely to survive during CRISPR-Cas9 editing^34^, so we investigated the expression of the different AS isoforms of TP53 in the SCR cells. Fig. S showed that the important isoforms, TP53-201, 209, 215, 219, 225 and 227, were all downregulated in SCR cells, whereas, TP53-204, 228 were upregulated (Supplementary Fig. 4A). The proteins of latter two isoforms are N-terminus truncated versions, which can compete with the functional p53 and inhibit apoptosis^35–38^. This result indicated that the AS of TP53 might have played a key role for the survival of SCR cells. We further looked at the expressions of some pro-apoptotic genes, such as FAS, FADD, BAX and CASP8, and found that they were all downregulated in various degrees (Supplementary Fig. 4B), suggesting that cells altered multiple pathways to deal with the stress brought by heavy DSBs.

### SCR reshaped the chromatin accessibility

On the one hand, epigenetics plays an important role in the regulation of gene expression, while on the other hand, we also wondered about the effect of changes in the primary structure of chromosomes on epigenetic modification. Therefore, we performed ATAC-seq for the SCR and NC cells. The overview of the peak distribution changes were shown in Fig. 2A and B. Here we further discuss the experimental results in more detail. The insert fragment size analysis indicates the distribution of nucleosomes on chromatins. NC showed a typical wave with a period of 200 bp, while the wave amplitude of SCR-L1 moderately decreased, and even more significantly, the wave of SCR-Alu was almost flat (Fig. 6A). This suggests that the nucleosome distribution in the SCR cells becomes less regular compared to the control cells. Presumably, the alteration in the nucleosome distribution would have broad impacts on the chromatin accessibility and the gene expression.

**Figure 6.**
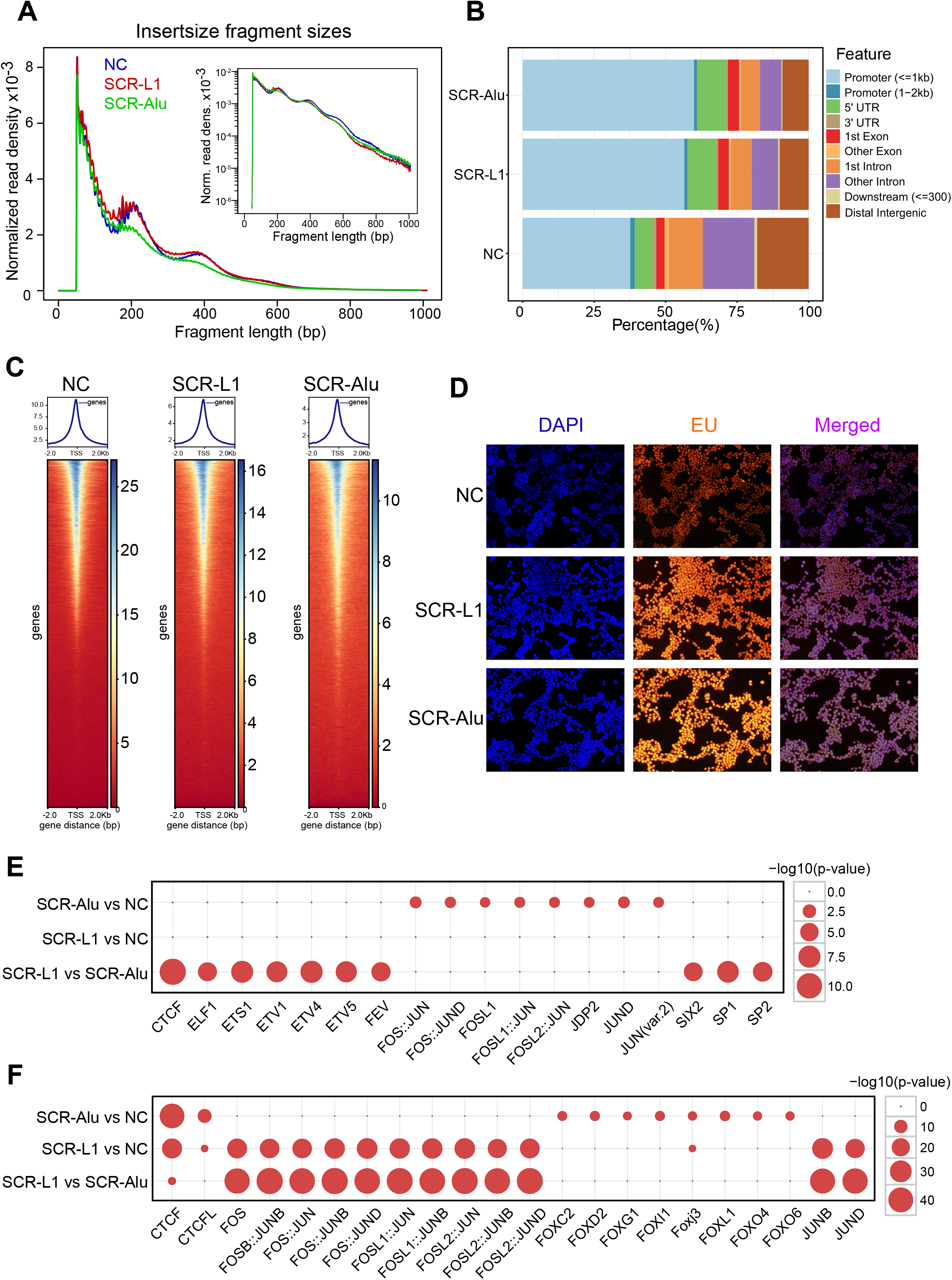
ATAC-seq revealed the change of chromatin accessibility of the SCR cells. A, the insert size distributions of the NC and SCR cells. B, The peak distributions in different genomic regions. C, the accessibilities flanking gene TSSs. D, EU staining to detect the new RNA synthesis. E & F, the bubble diagrams of significantly up-regulated motifs (E) and down-regulated motifs (F), the abscissa is the TF name, and the ordinate is the comparison between two samples. The size of the circle indicates the -log10 (P-value) of the corresponding TF motif.

The ATAC-seq data was also processed using Principal Component Analysis (PCA) (Supplementary Fig. 5A). The correlation analysis showed that SCR-L1 was closer to NC, while SCR-Alu was more different from those two groups (Supplementary Fig. B), which was consistent with the conclusion of transcriptome sequencing in Fig. 4 and Supplementary Fig. 1. The proportion of chromatin open regions in different positions, such as promoter, exon, intron, 5’UTR, 3’UTR, etc. also changed significantly. Only less than 40% of the open regions in NC are located in the promoter, while more than half of the open regions in SCR-L1 and SCR-Alu are located in the promoter regions (Fig. 6B). Moreover, statistics of the peak distribution flanking the gene transcription start sites (TSS) showed that nearly 80% of peaks distributed in the 1 kb window flanking the TSS in the SCR cells, but only ~50% peaks in the same window in NC cells (Supplementary Fig. 5C). These results suggested that the gene expression of cells after chromosome recombination might have become more active compared to control cells, though the total peak numbers dropped dramatically.

We then viewed the openness at the TSS. Nearly 170000 genes (including mRNA and lncRNA) were stacked together, and the heat map was drawn with ATAC peak values within 2 kb upstream and downstream of the TSS (Fig. 6C). Obviously, the diversity of the TSS openness is much larger in NC cells, whereas, the difference of openness between the high-open TSS and low-open TSS became smaller, and the open range near TSS of the high open genes become relatively wider in SCR cells. This result also indicated the disorder of the nucleosome distribution and gene-expression regulation.

Since ATAC-seq only measures the relative accessibility of chromatin and indicates the change of transcriptional activity, but not the actual RNA synthesis intensity, we performed the new RNA synthesis intensity assay in the three groups of cells. EU (5-ethynyl-2’uracil nucleoside) is a nucleoside analogue, which can be inserted into the newly formed RNA and emit fluorescence through conjugation reaction with fluorescent dye. Thus, the newly formed RNA can be detected by fluorescence microscopy or flow cytometry. Fig. 6D showed that the fluorescence increases from NC to SCR-L1 and then to SCR-Alu, which was what we had expected, because chromosome rearrangement inevitably led to a lot of wrong RNA products, and cells still need enough “right” RNA to ensure normal physiological function, which is bound to increase the total RNA. Therefore, the cells after SCR were in the state of active RNA synthesis and degradation.

To further analyze which transcription factors were related to the different peak distributions, motif analysis was conducted for the increased and decreased peaks respectively (Fig. 6E and F). Interestingly, CTCF was the top transcription factor related to the peak decrease in both SCR cells. CTCF is an insulator protein and is critical for maintaining the topologically associating domain (TAD) structure^39^. In the SCR cells, the chromatin accessibility of the CTCF binding sites was decreased, indicating that the binding between CTCF and chromosomal DNAs was blocked and the formation of TAD was impaired, which is consistent to recently reported that L1 and Alu repeats blueprint the chromatin macrostructure and TAD^40^. For the regions with increased peaks in SCR-Alu, JUND was the most remarkable transcription factor, which is also consistence to previously reported that the upstream regions of Alu tend to be associated with JunD^41^.

### The SCR cells acquired CRISPR-Cas9 resistance

Cas9 and sgRNA expression cassettes were integrated into the host genomes and forced to express in the presence of puromycin. However, the growth and proliferation of the SCR cells after recovery was similar to that of the NC cells, without significant apoptosis observed, which indicated that Cas9 was no longer continuously cleaving the chromosomal DNAs. Theoretically, Cas9 or sgRNA, or both of them, should have been inhibited. Since the Cas9 version used here has a Flag-tag on the C-terminus (the same version as in lentiCRISPR v2), we detected the expression of Cas9 in three groups of cells with anti-Flag antibody. Western blot showed that Cas9 proteins were almost undetectable in the two SCR cells in two independently screened cell pools respectively, whereas, the Cas9 proteins were highly expressed in the NC cells (Fig. 7A). We checked the abundances of Cas9 mRNA in the RNA-seq data. Since Cas9 and puromycin resistance gene (Puro^R^) were from the same plasmid and integrated into the genome by SB100X transposase simultaneously, the expression of Puro^R^ is an ideal internal reference for the expression of Cas9. We compared the RNA sequencing reads of the three groups of cells and found that despite the high expressions of Puro^R^ in all of them, the Cas9 mRNA was only highly expressed in the NC cells. However, in both of the two SCR cells, the Cas9 mRNAs were no longer complete, with a loss of the central region (Fig. 7B). According to the structure of Cas9 protein, the missing part mainly includes three domains, RecI, RuvC and HNH. RecI is the domain for sgRNA binding, while RuvC and HNH contain the endonuclease domain^43,44^. The defect of these domains would certainly destroy the activity of Cas9. The quantitative comparison of the RNA expression of this region (nt: 1123-3123; aa: 375-1041) also showed significant drops in the two SCR cells, especially in SCR-L1 (Fig. 7C). Therefore, the SCR cells might have developed certain mechanisms to silence the expression of Cas9 or both Cas9 and sgRNA.

**Figure 7.**
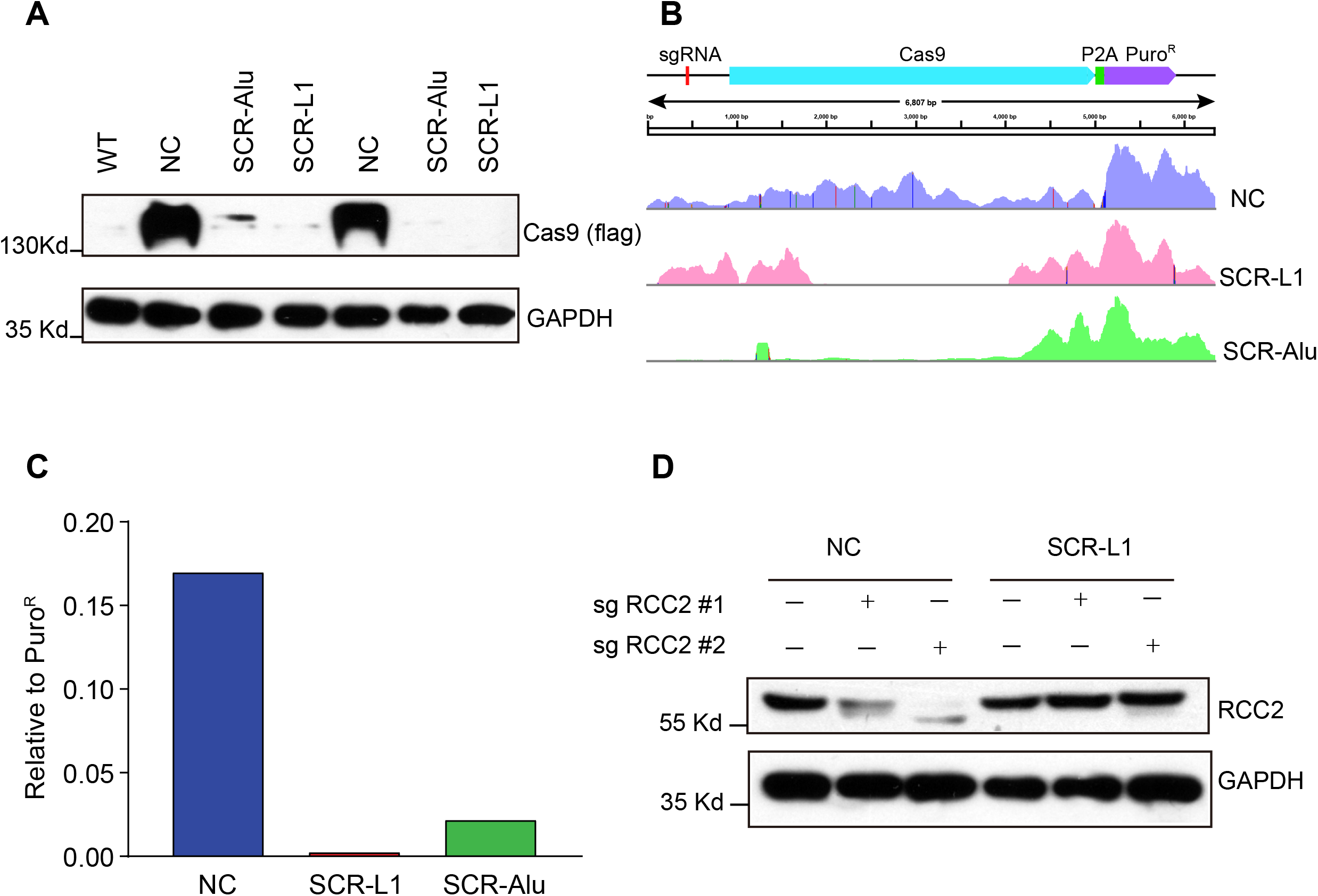
The CRISPR-resistance of the SCR cells. A, the expressions of Cas9 protein in NC and SCR cells. B, the RNA-seq read coverages of the sgRNA-Cas9-Puro^R^ cassette. C, the RNA-seq read coverage of the Cas9 (nt: 1123-3123) relative Puro^R^. D, the expressions of RCC2 protein after cells were transfected with pSB-sgRCC2-blast and selected by blasiticidin.

To test this hypothesis, we transfected the blasticidin resistance version of pSB-CRISPR with two sgRNA sequences aiming RCC2 gene into SCR-L1 cells. The knockout of RCC2 gene with pSB-CRISPR has been successfully performed previously, so it is a good positive control for testing whether the CRISPR-Cas9 system is working well. As what we have expected, Western blot showed that the RCC2 gene was successfully knocked out in the NC cells, whereas, RCC2 proteins were still highly expressed in SCR-L1 cells (Fig. 7D). This result showed that the SCR cells acquired CRISPR-Cas9 resistance by interfering with the Cas9 (or both Cas9 and sgRNA) expression (s).

Additionally, the Puro^R^ mRNA is in the same open reading frame with Cas9 mRNA, separated by P2A. After the central region of the mRNA was degraded, the translation of the 3’ part of the mRNA may start from one of the methionines at the Cas9 C-terminus, resulting in a Cas9 C-terminus fragment and an intact puromycin-N-acetyltransferase protein.

## Discussion

CRISPR genome editing tool, which was awarded the Nobel Prize in Chemistry in 2020^45^, is a powerful technology. In recent years, new applications based on it are also emerging rapidly and continuously. In this study, we developed a method, CReaC, which causes systematic chromosome rearrangement in the human genome, and deeply reshapes the landscapes of epigenetics and gene expression of the cells (Fig. 8). The wine yeast (*S. cerevisiae*) has a genome of only ~12 Mb, while the mammalian genomes are much larger and more complex, e.g., the haploid human and mouse genomes are ~3 Gb, so that it is impossible to generate SCR by synthesizing chromosomes at the current stage like the operation in yeast. CReaC provides an approach to study chromosome rearrangement in mammalian cells as well as in other complex eukaryotic cells.

**Figure 8.**
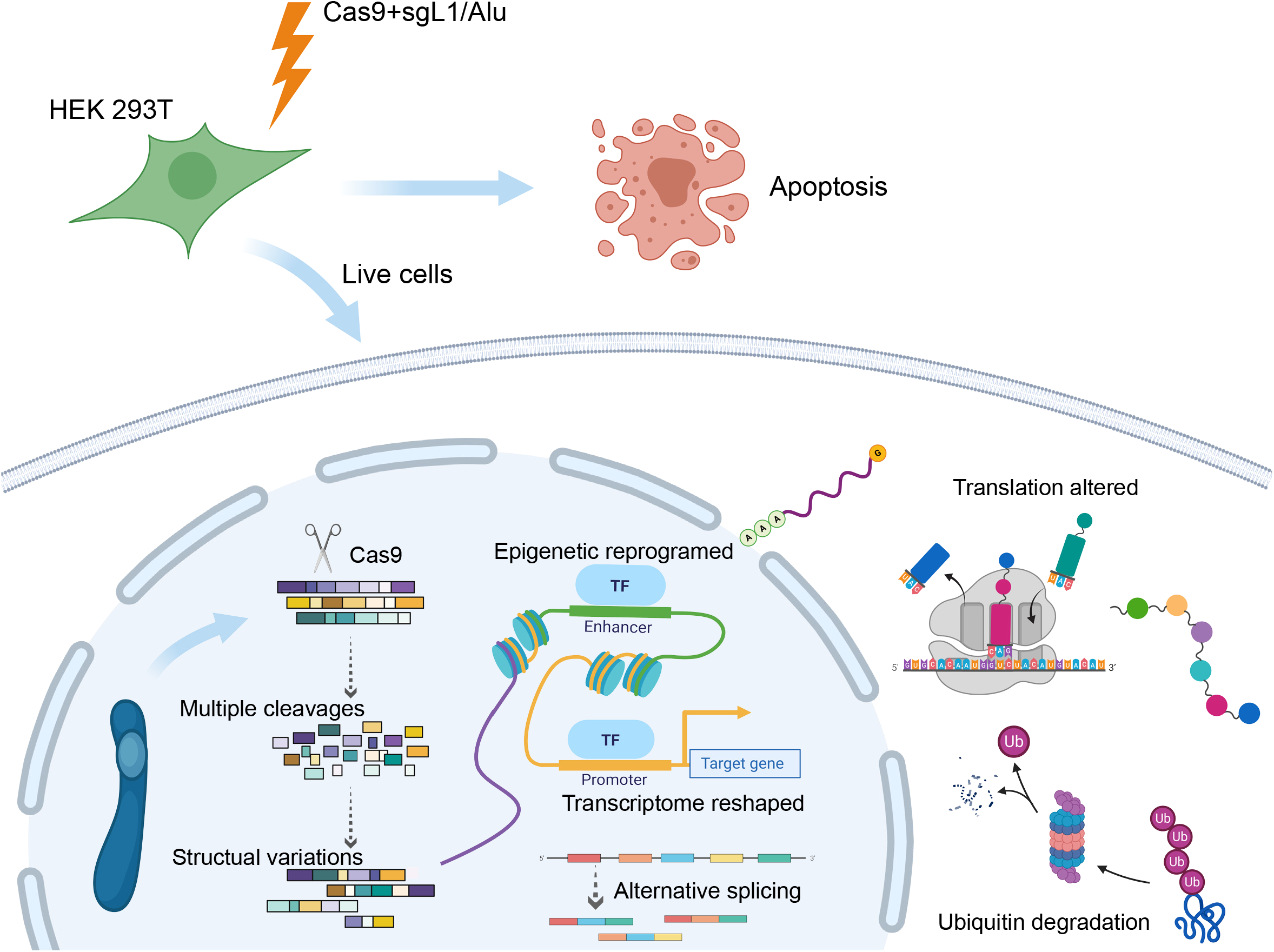
SCR brings great impact to cells. SCR is induced by targeting repetitive elements with CRISPR-Cas9. The 3D structure of the genome is changed; the distribution of nucleosomes becomes disorder; and the gene expression profile is reshaped.

In most cases, people use CRISPR-Cas9 to edit a specific site (two sites in diploid genomes) accurately, and try to avoid off-targets. However, in our study, Cas9 was directed to multiple sites by sgRNAs towards repetitive elements. The most famous multiple-site editing record were studies conducted by Yang and Church, in which more than 60 PERV genes were knocked out in porcine cells^46,47^, but whether cells can survive over hundreds, thousands or even more DSBs was yet unknown. Here we used sgRNAs with up to hundreds of thousands of matching sites to make DNA breakpoints and observed the subsequent changes, which, to our knowledge, is the first attempt so far. Although there have been several studies using CRISPR-Cas9 to generate chromosome rearrangements, those were only individual rearrangement instead of systematic rearrangement^48,49^.

The formation of the aneuploidy in carcinogenesis is a long-term and gradual process, and is difficult to investigate. Our research simulated this process in a considerably short time, which more intuitively explained the role of chromosome rearrangements in promoting cell carcinogenesis. Although the development and homeostasis of multicellular organisms are delicately tuned, our results suggest that their unicellular life mode, like immortalized cells or tumor cells, can be robust enough to tolerate severe chromosome rearrangements. The CNVs produced by chromosome rearrangements can only partially explain the changes in gene expression, and more changes were caused by gene expression regulations, in which epigenetics may play a key role. Our study showed that the distribution of nucleosomes tended to be disorder in the SCR cells. But it is unclear why the prime sequence changes in certain regions can cause the disorder distribution of nucleosomes globally.

It is interesting that SCR-Alu and SCR-L1 exhibit remarkably different, which might be due to that the copy number of Alu is far larger than that of L1 and the DNA DSBs in SCR-Alu are many more than those in SCR-L1, though the translocations and CNVs detected are similar. However, there is still other possibility that the cell status after SCR is not only determined by the chromosome rearrangement itself, but also is related to the functions of the elements where the DSBs take place. This result indicates that the physiological function of Alu might be even more important than L1 for its much higher copy number or for the transactivators interacting with them. Actually, recent studies showed that Alu elements are important for the formation of TADs and circRNAs^32,50,51^. Although the genomic fraction of Alu is smaller than that of L1 (11% vs 17%), it might be a more important composition than L1.

In addition, other repetitive elements, such as satellite DNAs, microsatellite DNAs, telomeres, centromeres or rDNAs can also be target options for CReaC and CReaC could be an approach for studying the functions of various repetitive elements.

While large-scale chromosome rearrangements may also be generated by using chemical or physical stimuli that cause DNA DSBs, our method has obvious advantages: the target site sequences and the potential number of target sites can be designed following requirements. Therefore, CReaC is more flexible and controllable than breaking DNA strands using chemical or physical reagents to generate random DSBs.

Currently, there have not been reports on CRISPR resistance. The concept of anti-CRISPR is defined as certain mechanisms that inhibit the function of CRISPR in prokaryotes^52,53^. Obviously, when exogenous Cas9 and sgRNA are introduced into eukaryotic cells, the cells will certainly have responses. Although CRISPR-Cas9 is a high-efficiency genome editing tool, it does exhibit low efficiency or even failure in many operations. It is reported that the causes of CRISPR failure are various and complicated^54^. Here we observed that the host cells may silence the Cas9 expression, especially the active center domain of Cas9 under certain circumstances, which provide a new angle to investigate the efficiency of CRISPR-editing. The study on the mechanism of CRISPR resistance may help promote or inhibit the activity of CRISPR-Cas9 in the future operations, and it can also improve the efficiency and safety of genome editing especially for therapeutic purpose.

Our current knowledge on the SCR cells is still limited, e.g., the structure variation of chromosomes and the gene expressions are all average exhibitions of cell pools. We are planning to perform WGS on single clones and single-cell transcriptome and ATAC-seq on the cell pools to study the epigenetic and transcriptomic changes at single-cell level. We are also interested in the 3D chromatin organization after SCR and can perform Hi-C assays to answer this question in the future.

## Materials and Methods

### Cells culture

The human immortalized normal renal cell line HEK 293T was obtained from the American Type Culture Collection (ATCC) and maintained in Dulbecco’s Modified Eagle Medium (DMEM, Gibco) supplemented with 10% fetal bovine serum (Gibco) and penicillin-streptomycin (HyClone) in a humidified incubator containing 5% CO_2_ at 37 °C.

### Plasmid construction, transfection and cell selection

The sgRNAs, sgL1 (TTCCAATCAATAGAAAAAGA) and sgAlu (TGTAATCCCAGCACTTTGGG) were designed according to the sequence of the most conserved regions of L1 and Alu respectively. The exact match number of these sgRNAs were searched against hg38 genome using a Perl script and confirmed using bowtie2 with the parameter of “no-1mm-upfront”^55^. The sgRNA sequences were cloned into pSB-CRISPR vector^17^, and a sgRNA sequence with no match in hg38 (CGCTTCCGCGGCCCGTTCAA) was used as negative control, the sgNC. Next, pSB-CRISPR and SB100X plasmids at a ratio of 10:1 were transfected into 293T cells using Lipofectamine 3000 (Invitrogen) following the manufacturer’s protocol. After 24 h the transfected cells were selected with puromycin at 1 μg/ml (Solarbio) for up to four weeks. The cells that survived were designated as SCR-L1, SCR-Alu and NC, respectively.

RCC2 knockout in SCR-L1 SCR-Alu and NC cells were also established using pSB-CRISPR system with sgRNA sequences, sgRCC2#1: TTGTGTCTGCAGCATGTGGGCGGANDsgRCC2#2: TGCAGTAGCAGCAGCGGCGG, as previously described^17^, and selected with Blasticidin S (Solarbio) at 10 μg/ml concentration for 3 weeks.

### Cell proliferation assay

Cell proliferation was measured using Cell Counting Kit-8 (CCK-8) assays. Briefly, 3 × 10^3^ SCR-L1, SCR-Alu and NC cells suspensions were seeded in a 96-well plates and were cultured for 24 h, 48 h and 72 h. A total of 10 μl of CCK-8 solution (APExBIO) was added to each well for 2 h-incubation at 37 °C at the same time every day, and then the absorbance at 450 nm was recorded using an enzyme immunoassay analyzer (TECAN Spark 10M).

### Survival rate after transfection

1 × 10^4^ 293Tcells suspensions were seeded in a 96-well plates for 24 h, and then pSB-CRISPR-sgNC, pSB-CRISPR-sgL1 and pSB-CRISPR-sgAlu plasmids were transfected together with SB100X plasmid into the cells using Lipofectamine 3000 (Invitrogen) following the manufacturer’s protocol. After 24 h the transfected cells were selected with puromycin at 1 μg/ml (Solarbio). CCK-8 assays as above were performed to detect cells survival rates at the 0 h, 24 h, 48 h, 72 h, 96 h, 120 h, 144 h, 168 h after selection.

### RNA synthesis detection

Cell-Light assay, based on combination of EU and Apollo fluorescent dyes, was used to detect cells RNA synthesis. 1 × 10^5^ SCR-L1, SCR-Alu and NC cell suspensions were seeded in 12-well plates for 24 h. Cells were fixed and stained for new synthesized RNA following the protocol of the Cell-Light EU Apollo567 RNA Imaging Kit (Ribobio), and then stained for DNA using DAPI. Finally, the images were observed and recorded using a fluorescence microscope (Olympus IX71, Tokyo, Japan).

### Cytogenetics analysis

Cytogenetics analysis of SCR-L1, SCR-Alu, NC were performed using G-banding techniques. Briefly, the cells were incubated with 0.06 μg/μL of colcemid for 2.5 h at 37°C, and then trypsinized, resuspended, centrifuged. The cells were then incubated in 0.075 M potassium chloride for 30 min at 37°C and fixed with Carnoy’s solution 3:1 (acetic acid:methanol). The metaphase chromosomes were analyzed for G-banding (500 band level) by Guangzhou LanGuang Co., Ltd. At least 5 cells were analyzed for each group of cells, and abnormality recognition and karyotype nomenclature were performed as the International System for Human Cytogenetic Nomenclature (ISCN).

### Western blot

Western blot were performed as previously described^56^. Briefly, the cells were collected and lysed with RIPA buffer (50 mM Tris-HCl [pH 8.0], 5 mM EDTA, 150 mM NaCl, and 0.5% Nonidet P-40 and aprotease and phosphatase inhibitor cocktail (Bimake, Shanghai, China). Equal amounts of protein were separated by 10% SDS-PAGE gel and transferred onto a polyvinylidene difluoride (PVDF) membranes (Millipore). The membranes were incubated with RCC2 (CST, #5104) and GAPDH (CST, #2118) antibodies overnight at 4 °C. Secondary antibodies (Transgen Biotech, #HS101) were incubated for 1 hour at room temperature. The membranes were visualized using ECL detection reagents (Beyotime #P0018A, Shanghai, China).

### Whole genome resequencing and Somatic variation analysis

Next generation sequencing library preparations were constructed following the manufacturer’s protocol (NEBNext® Ultra™ DNA Library Prep Kit for Illumina®). The libraries were sequenced using the Illumina HiSeq instrument with PE150 configuration by Genewiz Co., Ltd, Guangzhou, China. The clean data were aligned with human genome hg38 using BWA (version 0.7.12)^57^. CREST^58^ and Control-FREEC (version 10.6)^59^ were used to analyze the genomic structure variations.

### Whole transcriptome sequencing and analysis

Total RNA of the SCR-L1, SCR-Alu and NC cells was isolated using Trizol reagent (Invitrogen) according to the manufacturer’s instructions. RNA integrity and quantity were finally measured using RNA Nano 6000 Assay Kit (Agilent) of the Bioanalyzer 2100 system. The libraries were prepared and sequenced using Illumina sequencing by Novogene Co,. Ltd, Beijing, China. The raw reads were processed by removing the adaptor reads and low-quality tags. Clean reads for each sample were mapped to hg38 using the software HISAT2^60^. FPKM, the number of Fragments Per Kilobase of gene sequence per Millions base pairs sequenced was used to quantify the expression levels of a mRNA or lncRNA. The differential expression analysis of two conditions was performed using the edgeR R package (version 3.22.5). The P values were adjusted using the Benjamini & Hochberg method. Corrected P value < 0.05 and absolute foldchange > 2 were set as the threshold for significantly differential expression.

### ATAC-seq Library Preparation and Sequencing

ATAC-seq was performed according to the published protocol^61^. Briefly, when SCR-L1, SCR-Alu and NC cells were grown to 70~80% confluence, 5×10^5^ viable cells were lysed in 10 mM tris-HCl (pH 7.4), 10 mM NaCl, 3 mM MgCl_2_, and 0.1% (v/v) Igepal CA-630 and the nucleus was extracted. Transposition reaction was performed using TruePrep® DNA Library Prep Kit (Vazyme) at 37°C for 30 min, followed by purifing immediately. The libraries were amplified for 15 cycles using TruePrep® DNA Library Prep Kit (Vazyme), and sequenced using Illumina NovaSeqTM 6000 by Guangzhou Gene Denovo Biotechnology Co., Ltd., Guangzhou, China. After removing adapters and low quality reads, Bowtie2^55^ with the parameters –X2000 and –m1 was used to align the clean reads from each sample against hg38 genome assembly, and the reads aligned to the mitochondria were filtered. Peaks were called using MACS2 (version 2.1.2)^62^ with parameters “--nomodel --shift -100 --extsize 200 -B -q 0.05”. The DiffBind was used to analyse peak differences across groups, significant differential peaks were filter with FDR<0.05 in two comparison groups. Peak related genes were confirmed using ChIPseeker (version v1.16.1)^63^, besides with the distribution of peak on different genome regions (promoter, 5’UTR, 3’UTR, exon, intron, downstream and intergenic).

### The analysis on copy number variations

The genome was divided into equal intervals of 500 kb, and the differences of sequence read matches of WGS were calculated. The differences of matches per interval between SCR-L1 or SCR-Alu and NC represent the copy number variations (Fig. 2A and B; Fig. 3C). The regions with high CNVs in both SCR-L1 and SCR-Alu are the common high CNV regions. To analyze the translocation events in the common high CNV regions, each translation site were intersected with CNV coordinates using bedtools, the translocation/inversion sites located in the common high CNV regions were counted. For random regions, the regions with the same bin size of each CNV regions were shuffled 1000 times using bedtools, the amounts of translocation sites that these random regions were from each time were counted, and the distribution of translocation events were assessed (Fig. 3A and B). The genes in the common high CNV regions were annotated using Refseq annotation. Similarly, the TCGA clinical CNV data from Cosmic were also annotated using Refseq. The frequency of each genes observed in cosmic data were then calculated (Fig. 3E and F).

### Other bioinformatics analyses and charts

The Gene Ontology (GO), KEGG pathway, WikiPathway and Reactome enrichment analysis of differentially expressed genes was implemented by the clusterProfiler R package and CPDB online tools (cpdb.molgen.mpg.de/CPDB). Pathway terms with corrected P value < 0.05 were considered significantly enriched. The GO categories in biological networks were performed using Cytoscape plug-in, BinGo^64^. The bubble plots and the volcano plots were generated using ggplot package in RStudio. The heatmaps of H-cluster were generated using pheatmap R packages with hierarchical clustering method. The scatter plots (Fig. 2E and F; Supplementary Fig. 2) were generated using matplotlib package in Python. The Circos charts (Fig. 2A and B) were generated using RCircos package in RStudio. Briefly, the genome was divided into equal intervals of 500 kb, and the differences of sequence read matches of WGS, RNA-seq and ATAC-seq, and the peak coverages of ATAC-seq between SCR-L1 or SCR-Alu and NC were plotted as heatmaps or curves. The links at the center of charts were plotted according to the somatic structure variation data. Scripts for bioinformatics analyses were written in Perl, Python or R languages. The pictures of Fig. 1A and Fig. 8 were created with the aid of BioRender (biorender.com).

### Data access

The raw sequencing data were deposited in the Sequence Read Archive (SRA, https://www.ncbi.nlm.nih.gov/sra/) under accession numbers PRJNA732565 (WGS data), PRJNA731018 (RNA-seq data) and PRJNA731611 (ATAC-seq data), respectively.

## Supplementary Figure Legends

**Supplementary Figure 1.**
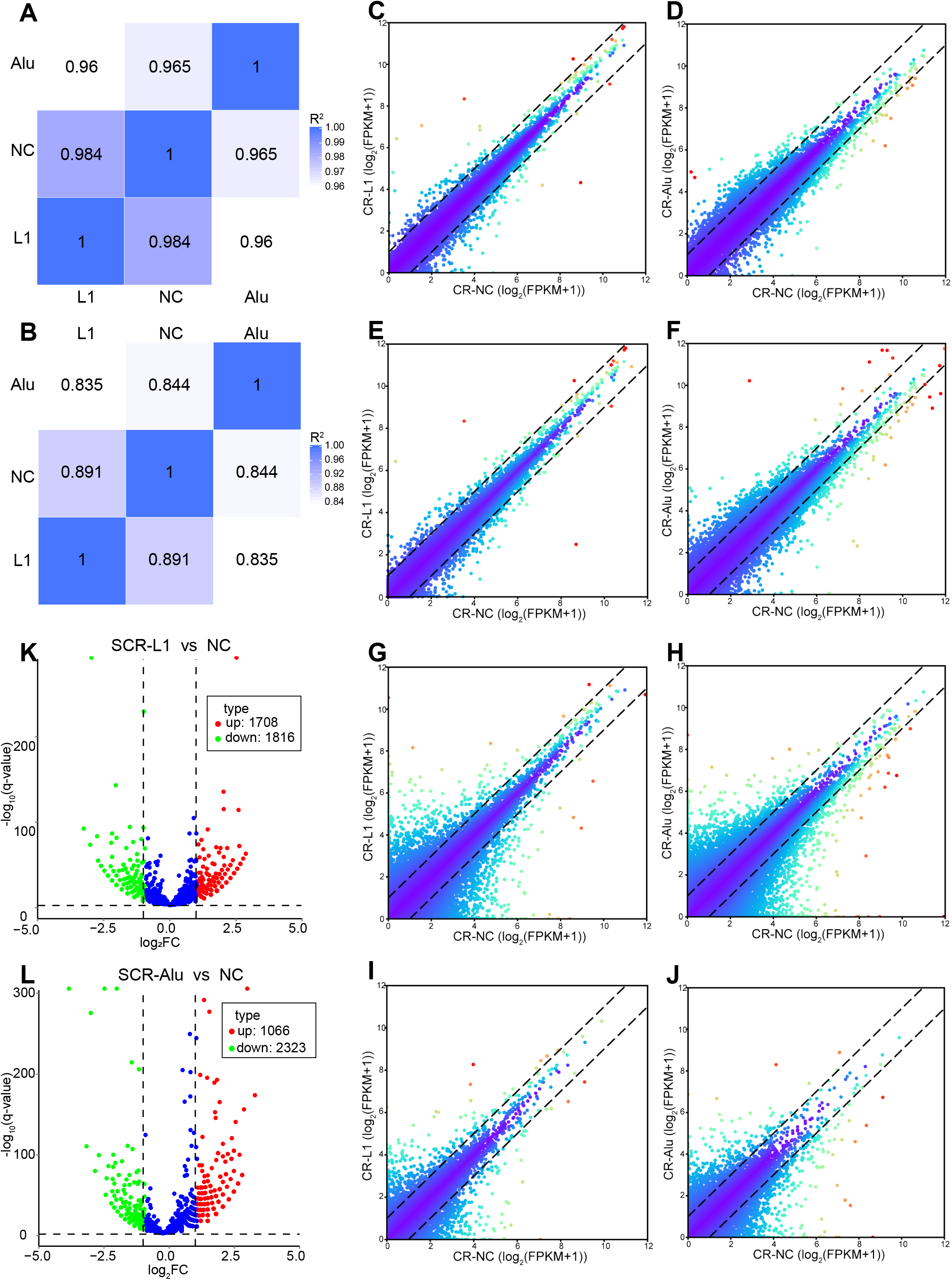
A-B, Pearson correlation between samples at gene level (A) and transcript level (B) in whole transcriptome sequencing. C-F, scatter plots show the differential expressions of mRNA (C & D) and lncRNA (E & F) at gene level in SCR-L1 vs NC (C & E) and SCR-Alu vs NC (D & F). G-J, scatter plots show the differential expressions of mRNA (J & H) and lncRNA (I & J) at transcript level in SCR-L1 vs NC (J & I) and SCR-Alu vs NC (H & J). K & L, volcano plots show the differential expressions of circRNA in SCR-L1 vs NC (K) and SCR-Alu vs NC (L).

**Supplementary Figure 2.**
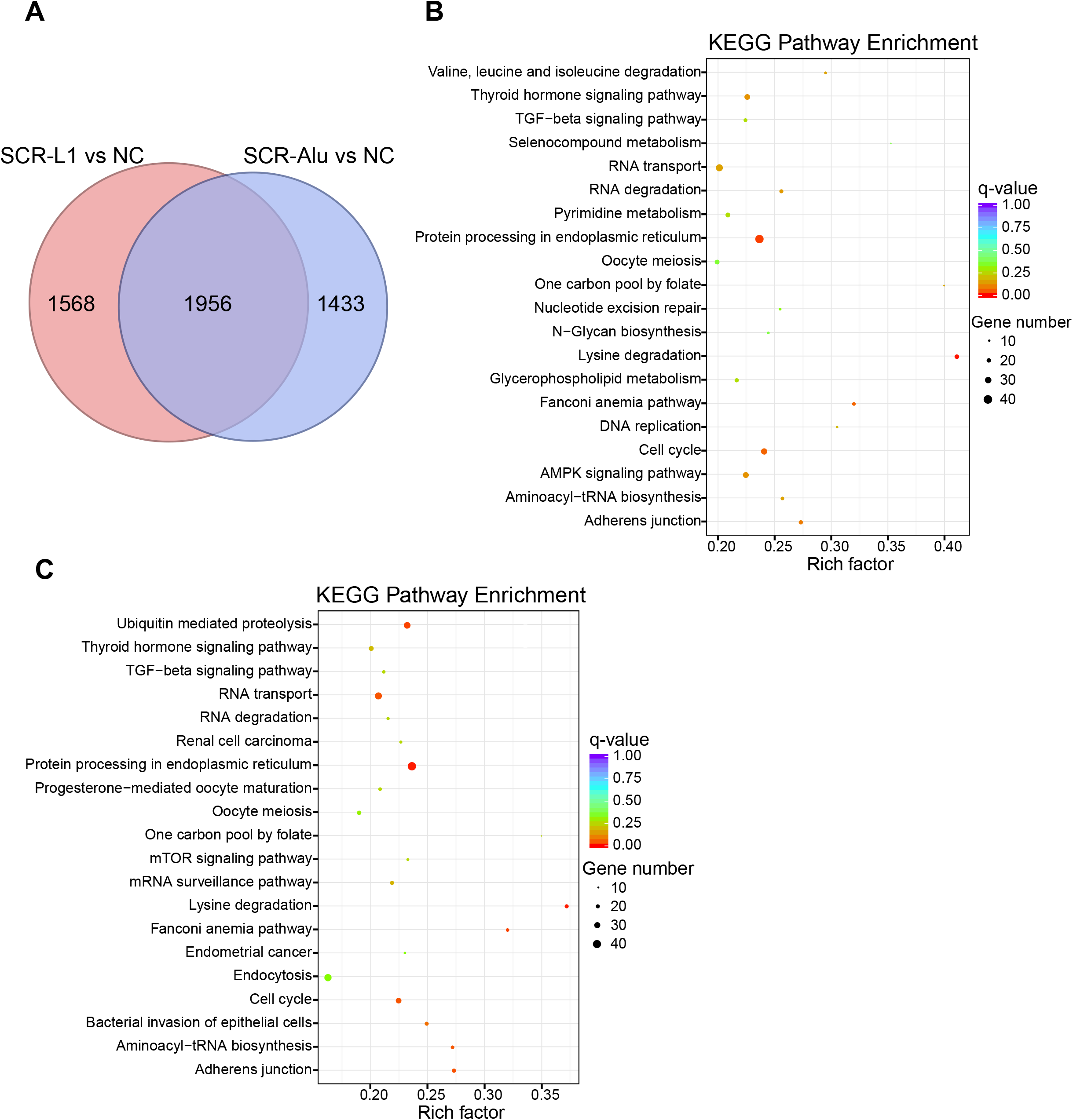
A, the numbers of the significant differential circRNA in SCR-L1 vs NC and SCR-Alu vs NC. B-C, the KEGG enrichment analysis of significant differential circRNA host genes of SCR-L1 vs NC (B) and SCR-Alu vs NC (C).

**Supplementary Figure 3.**
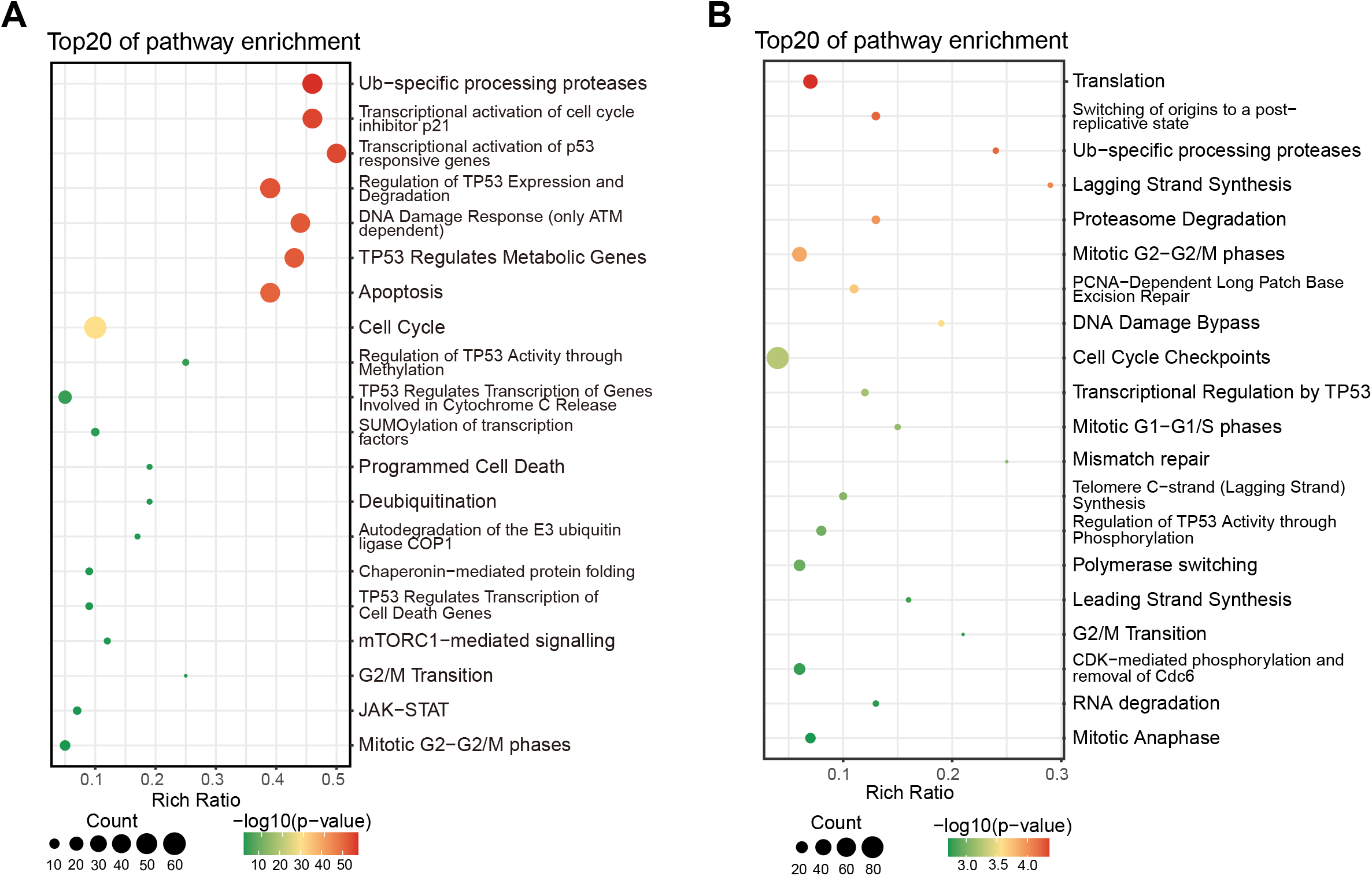
A, the 192 genes in panel 5B were analyzed using WikiPathway, Reactome and GO enrichments. B, the 520 genes in panel 5G were analyzed using WikiPathway, Reactome and GO enrichments.

**Supplementary Figure 4.**
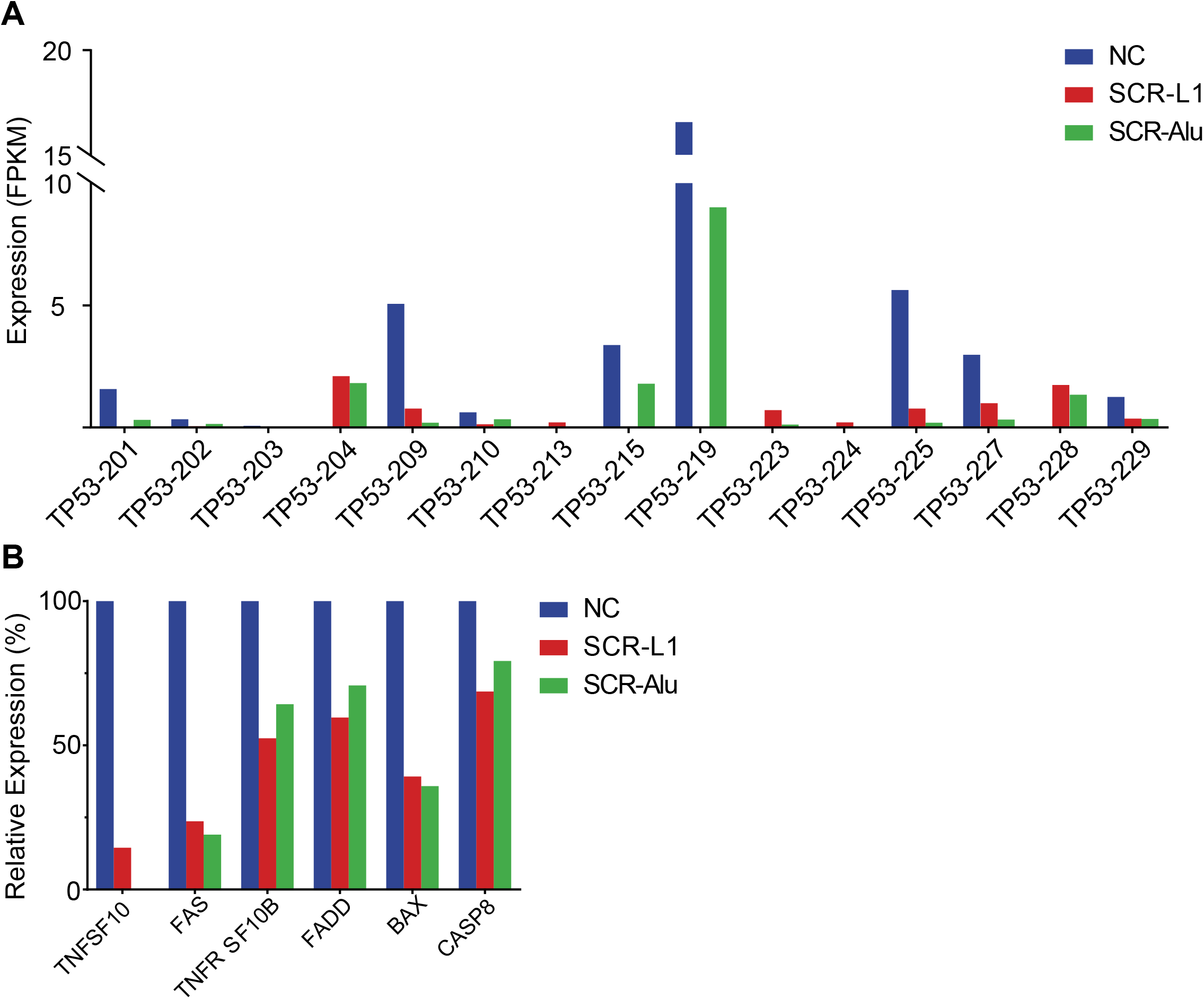
A, the alternative splicing of TP53 gene. B, the expression of some pro-apoptotic genes.

**Supplementary Figure 5.**
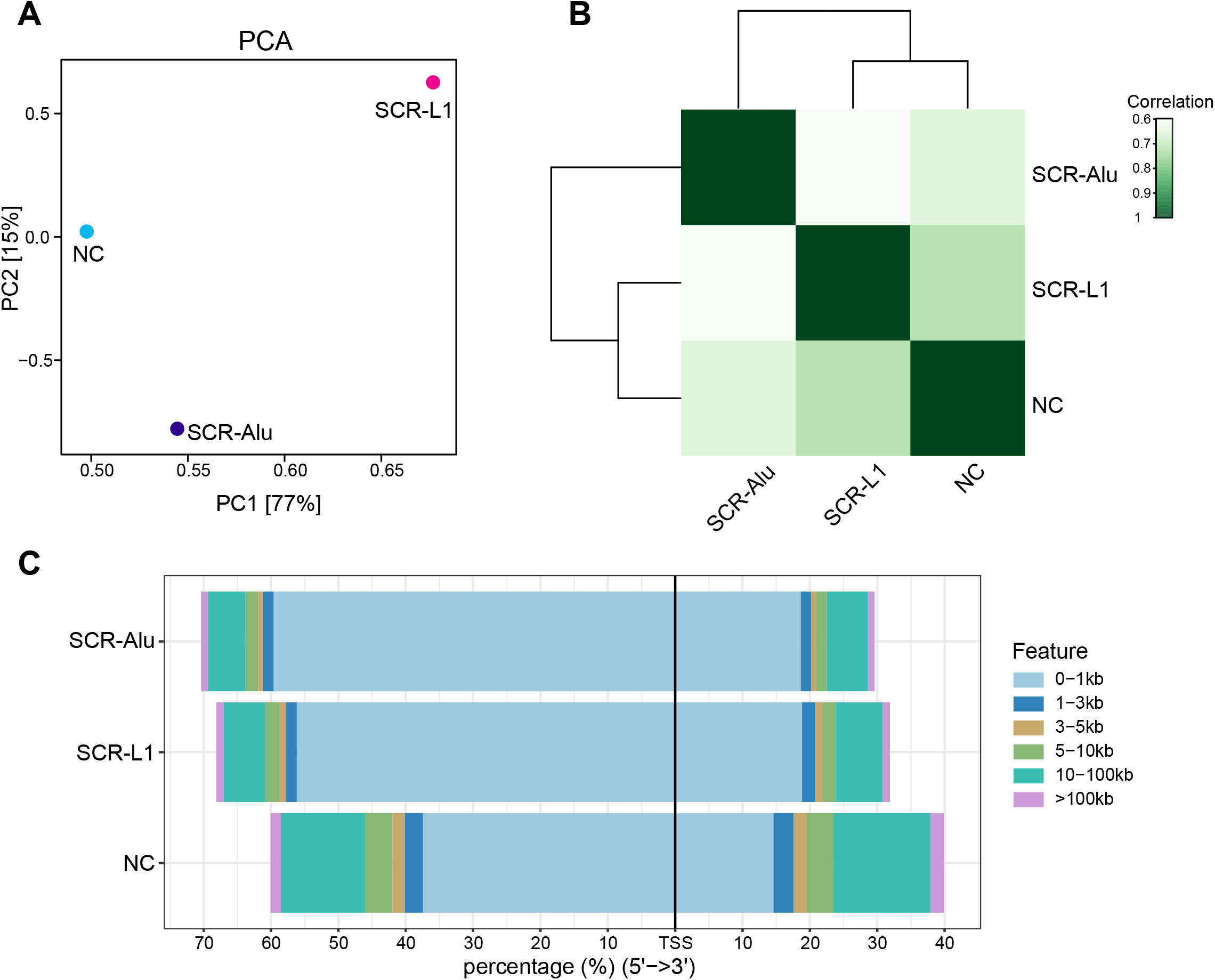
A, Principal component analysis (PCA) of ATAC-seq samples. PC1 and PC2 can explain 91.7% of the overall variance. B, Pearson correlation between ATAC-seq samples. C, distribution of distance to transcription start site (TSS) of common peaks in 3 samples. The abscissa indicates the percentage in different distances, and the ordinate indicates the sample.

## Notes

### Competing Interest Statement

The authors have declared no competing interest.

